# Lymphatic endothelial hemoglobin alpha promotes T cell accumulation and cardiomyocyte hypertrophy during cardiometabolic disease

**DOI:** 10.64898/2026.11.25.769328

**Authors:** Skylar A. Loeb, Wyatt J. Schug, Luke S. Dunaway, Clayton T. Grisius, Megan Burt, Zuzanna J. Juskiewicz, Madeline G. Jackson, Junjie Li, Shruthi Nyshadham, Olubodun M. Lateef, Juliana M. Parente, Laena Pernomian, Darla L. Tharp, Craig A. Emter, Camilla F. Wenceslau, Luis A. Martinez-Lemus, Antonio Abbate, Karen K. Hirschi, Miriam M. Cortese-Krott, Matthew J. Wolf, Brant E. Isakson

## Abstract

**Background:** Reduction in nitric oxide (NO) bioavailability is postulated to drive the progression of cardiometabolic diseases (CMDs), such as heart failure with preserved ejection fraction. The potent NO scavenger, hemoglobin alpha (Hbα), regulates endothelial-derived NO in health and disease. Whether lymphatic endothelial cells (LECs) express Hbα to modulate NO signaling and promote CMD is unknown.

**Methods:** We generated LEC-specific Hbα wildtype and knockout mice and fed them a high fat, high sucrose diet to investigate the role of LEC Hbα in CMD progression. To further define the necessity of LEC-derived NO in preventing CMD phenotypes, we generated LEC-specific endothelial nitric oxide synthase (eNOS) wildtype and knockout mice and subjected them to normal chow feeding.

**Results:** We found genes encoding Hbα were upregulated in the heart during CMD and that Hbα was enriched in cardiac lymphatic vessels. Genetic deletion of Hbα from LECs prevented CMD-induced cardiomyocyte hypertrophy and reductions in cardiac lymphatic density. Notably, our data demonstrate this occurred independent of changes in the coronary blood vasculature and was driven by local, rather than systemic, changes in NO. Scavenging of NO *in vitro* increased LEC expression of immunoregulatory genes, aligning with Hbα-mediated regulation of immune cell, and more specifically T cell, presence in the heart observed in CMD. Mechanistic studies *in vitro* revealed modulation of NO around LECs did not impact T cell activation, but instead adhesion – findings which were validated *in vivo*. Deletion of eNOS from LECs in otherwise healthy mice recapitulated CMD phenotypes of reduced cardiac lymphatic density, cardiomyocyte hypertrophy, and T cell enrichment in the heart.

**Conclusions:** These findings demonstrate LEC Hbα regulates local NO signaling during CMD to promote cardiac hypertrophic remodeling and inflammation. Thus, disruption of Hbα function in LECs poses as a potential therapeutic approach in CMDs associated with reduced NO bioavailability.

## Introduction

Cardiometabolic diseases (CMDs), in which chronic metabolic derangement is coincident with cardiovascular disease, are the leading cause of death worldwide.^1^ Within this broad category, heart failure with preserved ejection fraction (HFpEF) is a prime example, as metabolic comorbidities are strongly associated with HFpEF and are considered to drive its progression.^2^ While often heterogenous in clinical presentation, HFpEF is identified based on signs and symptoms of heart failure (e.g., dyspnea and/or exercise intolerance), a left ventricular ejection fraction ≥ 50%, and objective evidence of elevated left ventricle (LV) filling pressures.^3^ Over half of HFpEF patients also present with hypertrophy and/or concentric remodeling of the LV, with the former being associated with increased risk of heart failure-related death or hospitalization.^4^ While current therapies, such as sodium-glucose cotransporter 2 inhibitors and glucagon-like peptide-1 agonists, have improved patient quality of life and reduced worsening heart failure events or hospitalization, it remains crucial to define the mechanisms underlying cardiac hypertrophic remodeling in CMDs to identify novel therapeutic targets.^5, 6^

The current paradigm for HFpEF etiology is that cardiometabolic comorbidities drive a systemic pro-inflammatory and pro-oxidative state, resulting in reduced nitric oxide (NO) bioavailability.^2^ Decreased NO bioavailability has been well-documented in both HFpEF animal models and patients.^7–12^ Systemically, this can promote endothelial dysfunction and impaired vascular reserve.^13^ Locally in the heart, reduced NO signaling directly disturbs cardiomyocyte homeostasis. It has been shown in both *in vitro* and *in vivo* models of cardiac hypertrophy that NO signaling prevents cardiomyocyte hypertrophic remodeling.^14, 15^ Thus, decreased NO bioavailability links cardiometabolic comorbidities to the downstream HFpEF phenotype.

While other cell types of the heart (i.e., blood endothelial cells [BECs], cardiomyocytes, and fibroblasts) have been more extensively characterized in the context of CMDs, such as HFpEF, lymphatic endothelial cells (LECs) remain largely understudied. The cardiac lymphatic system plays crucial roles in maintaining fluid homeostasis, clearing waste, and regulating immune responses in the heart. Like BECs, LECs rely on balanced NO levels for proper function.^16^ Multiple lines of evidence have shown lymphatic dysfunction occurs in metabolic diseases when NO bioavailability is dysregulated.^17, 18^ However, little evidence exists investigating these mechanisms in the heart. It has been demonstrated that HFpEF patients have reduced cardiac lymphatic density and that stimulation of lymphangiogenesis in murine HFpEF prevented inflammation and partially prevented cardiac hypertrophy.^19^ This finding suggested lymphatic vessels are key contributors to cardiac inflammation during HFpEF. Recent work implicated T cell-mediated inflammation as a driver of HFpEF. Human HFpEF hearts were revealed to be enriched in activated T cells.^20^ *In vivo* studies also revealed that accumulation of T cells in the heart promotes cardiomyocyte hypertrophy.^21, 22^ While reduced lymphatic density and T cell-mediated inflammation have both been independently shown to drive CMD progression, no work has yet proven a connection between these two phenomena.

We have shown the potent NO scavenger, hemoglobin alpha (Hbα), is expressed in resistance artery endothelial cells.^23–26^ There, it interacts with endothelial nitric oxide synthase (eNOS) to scavenge eNOS-derived NO and regulate vascular tone.^25, 27, 28^ To date, no evidence has shown LEC expression of Hbα. Though, recent work from our group has shown Hbα expression is dysregulated in BECs during disease (i.e., iron deficiency anemia) to modulate NO levels.^29^ As NO bioavailability is reduced during HFpEF and lymphatic vessels have been linked to driving the phenotypes associated with reduced NO bioavailability, we hypothesized LECs express Hbα during CMD to reduce NO signaling in the heart, leading to cardiac inflammation and subsequent cardiomyocyte hypertrophy.

## Methods

### Data Availability

Newly generated RNA-sequencing data are deposited in NCBI’s Gene Expression Omnibus repository and are accessible through the GEO accession numbers GSE355124 and GSE355129. All other data supporting the findings of this study are available from the corresponding author upon reasonable request. Please see the Major Resources Table and Extended Materials and Methods in the Supplementary Materials.

## Results

### Hemoglobin alpha is enriched in cardiac LECs during CMD

Many lines of evidence have shown NO bioavailability is reduced in CMDs.^7–12^ While studies suggest scavenging by excess reactive oxygen species as the driver of this phenomenon, antioxidant therapy has remained unsuccessful in the treatment of cardiovascular diseases.^30^ To determine an alternative mechanism through which NO signaling may be dysregulated during CMD, we performed bulk RNA-sequencing on heart tissue from mice subjected to 15 weeks of high fat diet (60% kcal fat) plus L-NAME (0.5 g/L in drinking water) to model HFpEF.^31^ We identified *Hba1*, encoding the potent NO scavenger Hbα, was upregulated in murine HFpEF hearts (**Figure 1A**). Publicly available bulk RNA-sequencing^32^ from human control, HFpEF, and heart failure with reduced ejection fraction (HFrEF) hearts similarly showed *HBA1* and *HBA2* were up-regulated in both HFpEF and HFrEF hearts, though to a much lesser extent in HFrEF (**Figure 1B, Supplementary Figure 1A**). Of note, the genes encoding Hbα were the most highly upregulated genes in human HFpEF, while other red blood cell genes (*ALAS2, GATA1, MFRN1, GYPA*) were undetected, suggesting a non-erythroid source of *HBA1/2* up-regulation. Publicly available single-cell RNA-sequencing of murine HFpEF and control hearts^33^ revealed *Hba1* had the highest average expression in pan-endothelial cells and that *Hba1* is up-regulated in pan-endothelial cells during HFpEF (**Supplementary Figures 1B-1C**). Publicly available single-nucleus RNA-sequencing from human HFpEF hearts^20^ revealed this pan-endothelial up-regulation observed in the mouse may be in part derived from LECs, which showed an increase in *HBA2* transcripts in HFpEF patients (*HBA1* was undetected in LECs) (**Figure 1C**). To validate this finding *in vivo*, we placed mice on an 18-week-long high fat, high sucrose (HFHS; 40% kcal each) diet to model CMD (**Figure 1D**). While this model does not recapitulate all features of HFpEF, it produces many of the comorbid phenotypes considered to drive HFpEF (e.g., obesity, glucose intolerance, and coronary vascular dysfunction), making it a good model for studying early disease-driving mechanisms.^34, 35^ Importantly, this model also allows us to study CMD without confounding exogenous manipulation of NO bioavailability with L-NAME as occurs in the murine HFpEF model. Immunostaining on the LVs from HFHS-fed mice and age-matched, normal chow (NC)-fed controls revealed Hbα is enriched in both BECs and LECs during CMD (**Figures 1E-1F**). Of note, Hbα enrichment in LECs was five-fold greater than that of BECs (**Figure 1G**). Taken together, these data identify LECs as a major source of increased Hbα during CMD.

**Figure 1.**
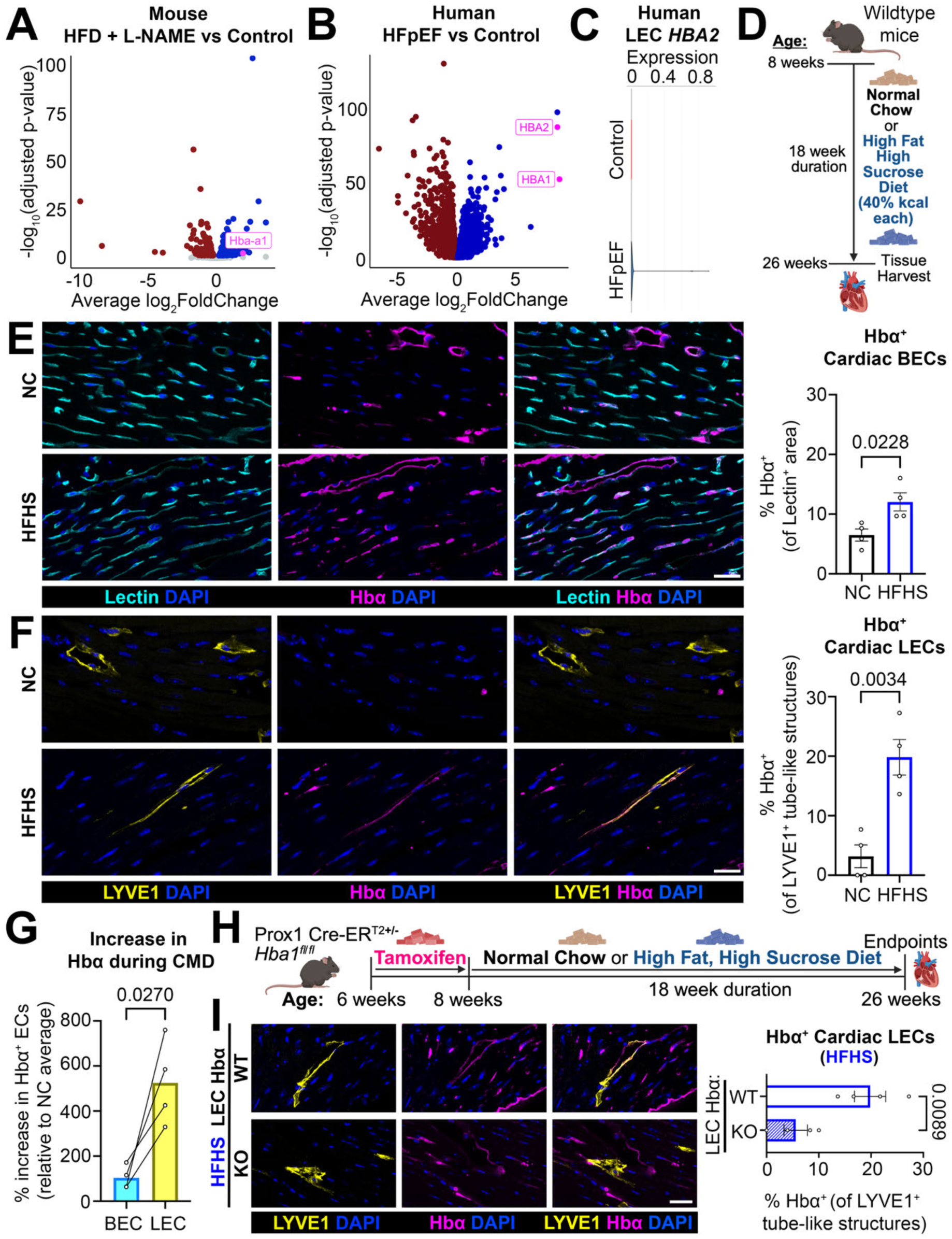
Hemoglobin alpha is upregulated in cardiac lymphatic endothelium during cardiometabolic disease. (A-B) Differentially expressed genes from bulk RNA-sequencing of heart failure with preserved ejection fraction (HFpEF) versus control hearts derived from (A) mice and (B) humans. Mouse model = 15 weeks of high fat diet (HFD) and L-NAME (0.5 g/L in drinking water). Statistical analysis for mouse data performed with DESeq2. Human data re-analyzed from *Hahn et al*.^32^ (C) *HBA2* expression in cardiac lymphatic endothelial cells (LECs) from control donors and HFpEF patients. Data re-analyzed from *Hahn et al*.^20^ (D) Experimental scheme. Wildtype (WT) mice were fed a high fat, high sucrose (HFHS) diet for 18 weeks to model cardiometabolic disease (CMD). Control mice were fed normal chow (NC). (E-F) Representative immunostains of left ventricles from NC- and HFHS-fed mice for hemoglobin alpha (Hbα) and either (E) lectin or (F) LYVE1. Percent of Hbα-expressing (E) blood endothelial cells (BECs) or (F) LECs quantified to the right. N=4 mice/group. Scale = 25 μm. (G) Percent increase in Hbα-expressing BECs and LECs during CMD. N=4 mice/group. (H) Experimental scheme for Prox1 Cre-ER^T2+/-^ *Hba1^fl/fl^* (LEC Hbα-WT and -knockout [KO]) mice. (I) Representative images of left ventricles from HFHS-fed LEC Hbα-WT and -KO mice stained for Hbα and LYVE1, with quantification of percent of LECs expressing Hbα. HFHS WT values from F. N=4 mice/group. Scale = 25 μm. Statistical significance determined by (E, F, I) unpaired t-test or (G) paired t test.

### LEC Hbα regulates structural changes in the heart during CMD

To investigate the role of LEC Hbα in CMD progression, we generated *Prox1 Cre-ER^T2+/-^ Hba1^fl/fl^* (LEC Hbα-wildtype [WT] and -knockout [KO]) mice to selectively delete Hbα from LECs. Six-week-old male and female mice were administered tamoxifen to induce the genetic deletion (validated by both DNA excision from ear tissue and a reduction in Hbα-expressing cardiac lymphatic vessels) prior to undergoing 18 weeks of NC or HFHS feeding (**Figures 1H-1I, Supplementary Figure 2A**). Deletion of LEC Hbα did not impact BEC expression of Hbα (**Supplementary Figure 2B**). There is evidence that *Prox1* is expressed in cardiomyocytes during development.^36^ To ensure LEC specificity in our results, we generated a cardiomyocyte-specific Hbα-KO mouse (*Tnnt2^MerCreMe/+^Hba1^fl/fl^)*, from which we observed successful DNA excision from heart tissue (**Supplementary Figures 2C-2D**). However, we did not observe changes in cardiac Hbα levels in cardiomyocyte Hbα-KO mice nor could we visualize Hbα in cardiomyocytes of cardiomyocyte Hbα-WT hearts by immunostaining, strongly suggesting that Hbα is not expressed in cardiomyocytes (**Supplementary Figures 2E-2F**). To further validate the LEC specificity of any major findings, we generated a second LEC Hbα-KO mouse using a *Flt4-CreER^T^*^2^, which we validated by both DNA excision from ear tissue and a visualized loss of Hbα immunofluorescent signal in cardiac LECs of HFHS-fed KO mice (**Supplementary Figures 2G-2I**).

We found male HFHS-fed mice had increased body mass, cholesterol levels, and high-density lipoprotein levels, all of which occurred independently of LEC Hbα expression (**Supplementary Figures 3A-3C**). Loss of Hbα from LECs also had no effect on red blood cell count, blood hemoglobin levels, mean arterial pressure, or heart rate. (**Supplementary Figures 3D-3G**). Echocardiography revealed that while ejection fraction was preserved amongst all groups, male HFHS-fed LEC Hbα-WT mice had reductions in stroke volume, end diastolic volume, and end systolic volume, as well as increased LV wall thickness. These phenotypes were all rescued in the absence of LEC Hbα (**Figures 2A-2E**). Further, hearts from male HFHS-fed mice also weighed more only when LEC Hbα was present (**Figure 2F**).

**Figure 2.**
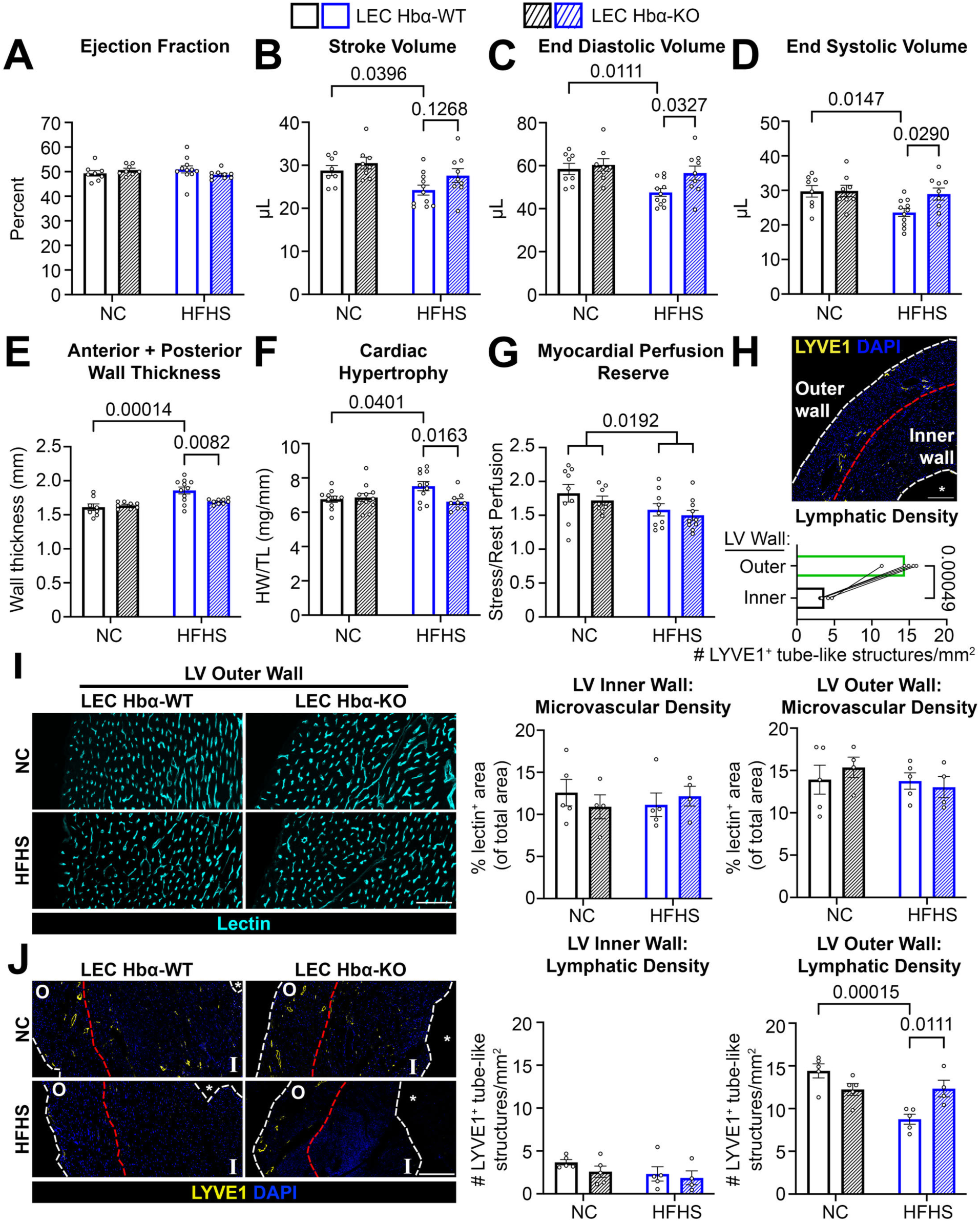
LEC Hbα regulates cardiac structure during CMD independent of changes in the coronary blood vasculature. (A-E) (A) Left ventricular ejection fraction, (B) stroke volume, (C) end diastolic volume, (D) end systolic volume, and (E) wall thickness measured from normal chow (NC)-fed and high fat, high sucrose (HFHS)-fed lymphatic endothelial cell (LEC) Hbα-wildtype (WT) and -knockout (KO) mice by echocardiography. NC LEC Hbα-WT and -KO groups: N=8 mice each; HFHS LEC Hbα-WT: N=11; HFHS LEC Hbα-KO: N=9. (F) Heart weight (HW) normalized to tibial length (TL) from all groups. NC LEC Hbα-WT: N=10 mice; NC LEC Hbα-KO: N=11; HFHS LEC Hbα-WT: N=11; HFHS LEC Hbα-KO: N=9. (G) Myocardial perfusion reserve measured via cardiac magnetic resonance imaging as a ratio of perfusion at stress (adenosine exposure) to perfusion at rest. NC LEC Hbα-WT: N=9 mice; NC LEC Hbα-KO: N=8; HFHS LEC Hbα-WT: N=9; HFHS LEC Hbα-KO: N=10. (H) Representative LYVE1 immunostain on a NC LEC Hbα-WT left ventricle (LV), with wall-specific quantifications of lymphatic density. N=5 mice. (I) Representative lectin staining on the LV outer wall from all groups, with wall-specific quantifications of microvascular density. NC and HFHS LEC Hbα-WT groups: N=5 mice each; NC and HFHS LEC Hbα-KO groups: N=4 each. Scale = 50 μm. (J) Representative LYVE1 immunostains on LVs from all groups, with wall-specific quantifications of lymphatic density. NC LEC Hbα-WT values from H. NC LEC Hbα-WT and -KO groups: N=5 mice each; HFHS LEC Hbα-WT: N=5; HFHS LEC Hbα-KO: N=4. For lymphatic density measurements (H, J): * denotes LV chamber. White dashed lines indicate LV wall borders; red dashed line indicates the approximate end of where lymphatics populate. Scale = 200 μm. Statistical significance determined by (A-G, I-J) 2-way ANOVA with Šídák’s multiple comparisons test or (H) paired t test.

It has been described that endothelial dysfunction often precedes the structural and functional changes occurring in the heart during CMD.^2, 37^ Upon examination of coronary vascular function using cardiac MRI, we found the vascular response to adenosine-induced stress was impaired in male mice with CMD, though this was not regulated by LEC Hbα (**Figure 2G**). The murine cardiac lymphatic system is highly localized to the outer 300 μm of the LV wall^38^ (**Figure 2H**), so to account for potential localized effects of LEC Hbα on the blood vasculature, we assessed microvascular density in the LV inner and outer walls separately. However, microvascular density was unchanged across all conditions in both regions of the LV wall (**Figure 2I, Supplementary Figure 4**). We did instead observe a reduction in cardiac lymphatic density only in the LV outer wall of male HFHS-fed LEC Hbα-WT mice, which was rescued in LEC Hbα-KO mice (**Figure 2J**). Of note, HFHS-fed female mice did not exhibit any changes in heart structure and function measured by echocardiography nor was cardiac lymphatic density altered (**Supplementary Figures 5A-5F**). Male mice were therefore used for the remainder of the study. These data together suggest LEC Hbα regulates structural changes in the heart during CMD and that these changes occur independent of the blood vasculature.

### LEC Hbα promotes cardiomyocyte hypertrophy during CMD through local reduction of NO signaling in the heart

To determine whether LEC Hbα promotes cardiomyocyte hypertrophic remodeling to regulate cardiac structure, cardiomyocyte cross-sectional area was assessed across the LV wall. Cross-sectional areas of cardiomyocytes from the inner wall were unchanged, even during CMD (**Figure 3A, Supplementary Figure 6A**). Conversely, cardiomyocytes localized to the LV outer wall became hypertrophic during CMD, and this phenotype was reliant on the expression of LEC Hbα (**Figure 3A**). We observed the same effect with *Flt4*-driven deletion of Hbα from LECs (**Supplementary Figures 6B-6D**). Cardiomyocyte-specific deletion of Hbα did not prevent cardiomyocyte hypertrophy in HFHS-fed mice, further pointing to LEC specificity of this phenotype (**Supplementary Figures 6E-6G**).

**Figure 3.**
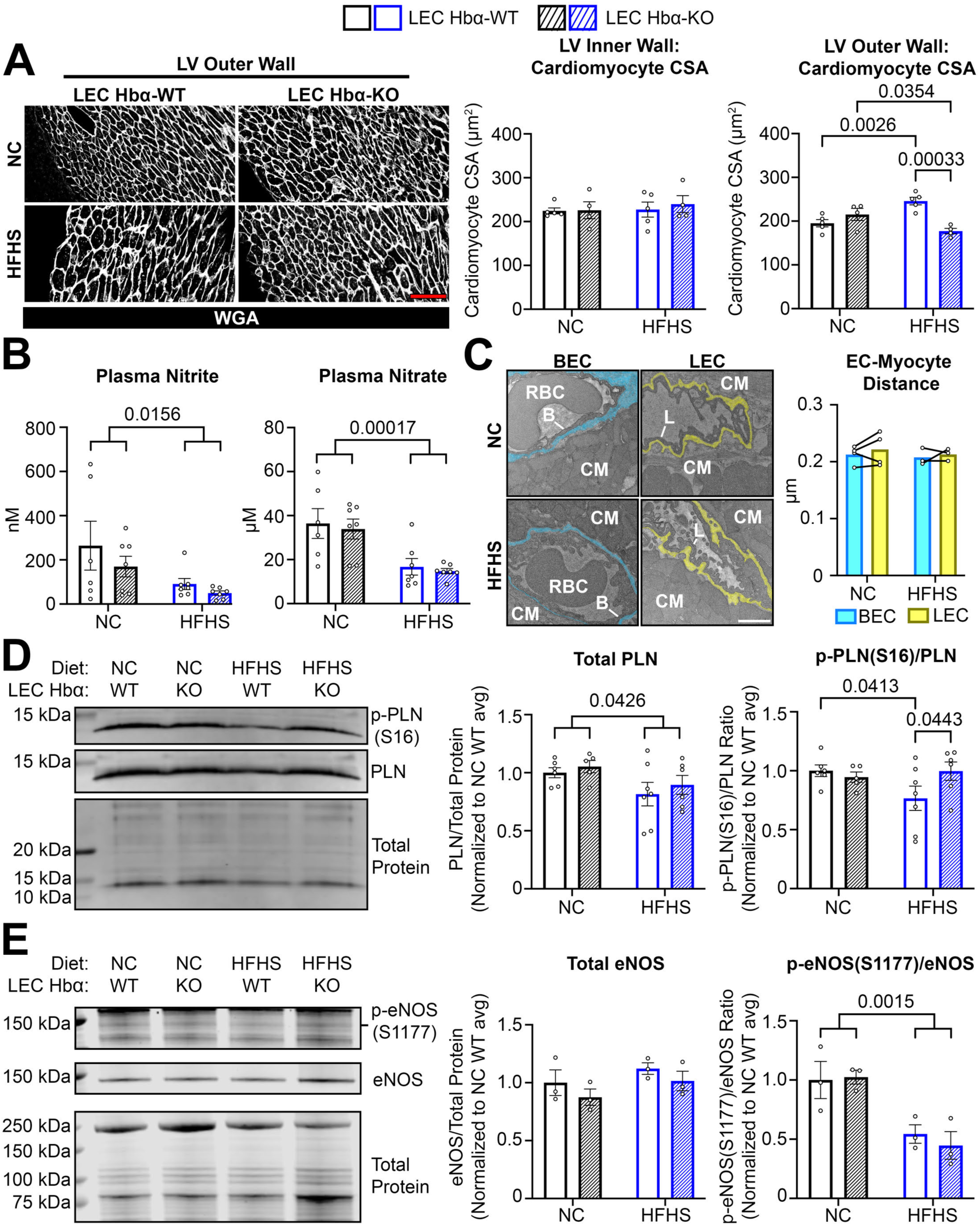
LEC Hbα promotes cardiomyocyte hypertrophy during CMD by locally reducing nitric oxide signaling in the heart. (A) Representative wheat germ agglutinin (WGA) stains from the left ventricle (LV) outer wall of normal chow (NC)-fed and high fat, high sucrose (HFHS)-fed lymphatic endothelial cell (LEC) Hbα-wildtype (WT) and - knockout (KO) mice, with wall-specific quantifications of cardiomyocyte cross-sectional area (CSA). NC and HFHS LEC Hbα-WT groups: N=5 mice each; NC and HFHS LEC Hbα-KO groups: N=4 each. Scale = 50 μm. (B) Plasma nitrite and nitrate levels. NC LEC Hbα-WT: N=6 mice. All other groups: N=7 each. (C) Representative transmission electron microscopy images from the heart showing blood and lymphatic capillaries and their neighboring cardiomyocytes with quantifications of blood endothelial cell (BEC) and LEC distance from the nearest cardiomyocyte. N=4 NC-fed LEC Hbα-WT mice, 3 HFHS-fed LEC Hbα-WT mice. Six distances were averaged per vessel, with 5 blood vessels and 5 lymphatic vessels separately averaged per mouse. Shaded regions (blue or yellow) indicate intercellular regions. RBC = red blood cell, B = blood endothelial cell, L = lymphatic endothelial cell, CM = cardiomyocyte. Scale = 1μm. (D) S16 phosphorylation and total phospholamban (PLN) from LV lysates, assessed by western blot. NC LEC Hbα-WT: N=6 mice; NC LEC Hbα-KO: N=5; HFHS LEC Hbα-WT: N=7; HFHS LEC Hbα-KO:N=6. (E) S1177 phosphorylation and total endothelial nitric oxide synthase (eNOS) from LV lysates, assessed by western blot. N=3 mice/group. Statistical significance determined by (C) 2-way repeated measures ANOVA or (A-B, D-E) 2-way ANOVA with Šídák’s multiple comparisons test.

As a potent NO scavenger, Hbα may be driving the described phenotypes through systemic and/or local regulation of NO levels. To assess systemic changes in NO bioavailability, we measured plasma levels of NO metabolites, nitrite and nitrate. Levels for both metabolites were reduced in HFHS-fed mice independent of genotype, suggesting that LEC Hbα does not promote the systemic reduction in NO bioavailability occurring in CMD (**Figure 3B**). This result is contrary to what was observed in pan-endothelial cell Hbα-KO mice (with *Cdh5*-driven deletion of Hbα from all endothelial cells), which had increased plasma nitrate levels and a trending increase in nitrite levels under normal conditions (**Supplementary Figure 7**).

We next examined whether LEC Hbα regulates NO signaling locally in the heart to mediate the whole-organ changes observed in mice with CMD. Transmission electron microscopy revealed capillary BECs and LECs are in equal proximity to their neighboring cardiomyocytes in both healthy and CMD mice (**Figure 3C**). Thus, LEC scavenging of NO via Hbα could directly impact NO signaling in neighboring cardiomyocytes. To determine whether LEC Hbα modulates NO signaling locally in the heart, we assessed the phosphorylation of phospholamban (PLN) at its serine 16 residue, which has been shown to be influenced by downstream NO signaling.^39, 40^ Immunoblotting revealed that while total PLN was reduced in HFHS-fed mice independent of LEC Hbα, the phosphorylation of PLN at serine 16 was reduced in HFHS-fed LEC Hbα-WT mice and subsequently restored in knockout mice (**Figure 3D**). Importantly, total eNOS protein levels were unchanged, aligning with the unaltered transcript levels for *Nos3*/*NOS3* observed in murine and human HFpEF (**Figure 3E, Supplementary Figures 8A-8B)**. eNOS phosphorylation at the serine 1177 residue was also reduced independent of LEC Hbα (**Figure 3E**). These data suggest that while eNOS activity may be decreased in the heart during CMD, it is not responsible for the changes in cardiomyocyte NO signaling observed. Thus, LEC Hbα promotes cardiomyocyte hypertrophy during CMD through local, and not systemic, regulation of NO bioavailability.

### LEC Hbα drives T cell accumulation in the heart during CMD

Dysregulation of NO levels has been shown to disturb lymphatic homeostasis.^17, 41^ To evaluate the potential consequences of Hbα upregulation in LECs, we mimicked its NO-scavenging function *in vitro* by treating human dermal LECs with an exogenous NO scavenger, PTIO (100 μM), for 24 hours and performed bulk RNA-sequencing (**Figures 4A-4B**). Through gene set enrichment analysis, we found pathways related to immune response and immunoregulation were activated in PTIO-treated LECs relative to control cells. (**Figure 4C**). To validate the potential role of Hbα-expressing lymphatic vessels in promoting inflammation, we analyzed cardiac immune cells using flow cytometry (**Supplementary Figure 9**). In line with the RNA-sequencing results, HFHS-fed LEC Hbα-WT hearts had an increased percentage of immune cells relative to NC-fed controls—an effect lost with the deletion of Hbα from LECs (**Figures 4D-4E**). We next analyzed macrophage, dendritic cell, and T cell populations in the heart, as these are the predominant immune cell types trafficked by lymphatic vessels during inflammation.^42^ While the percentage of macrophages and dendritic cells were both unchanged across all groups, the percentage of T cells was increased in HFHS-fed mice in an Hbα-dependent manner (**Figures 4F-4H, Supplementary Figure 10A**). *Flt4*-driven deletion of Hbα in mice with CMD achieved similar results (**Supplementary Figures 10B-10C**). Further phenotyping of cardiac T cells using flow cytometry revealed the percentages of both cytotoxic T cells and T helper cells were increased in HFHS-fed LEC Hbα-WT mice but not LEC Hbα-KO mice. (**Supplementary Figures 10D-10F**) Immunostaining for CD3^+^ T cells in the hearts of mice from all groups revealed the majority of T cells in the heart were localized in the LV outer wall and it was only in the outer wall where LEC Hbα regulation of T cell count was observed (**Figure 4I, Supplementary Figure 10G**).

**Figure 4.**
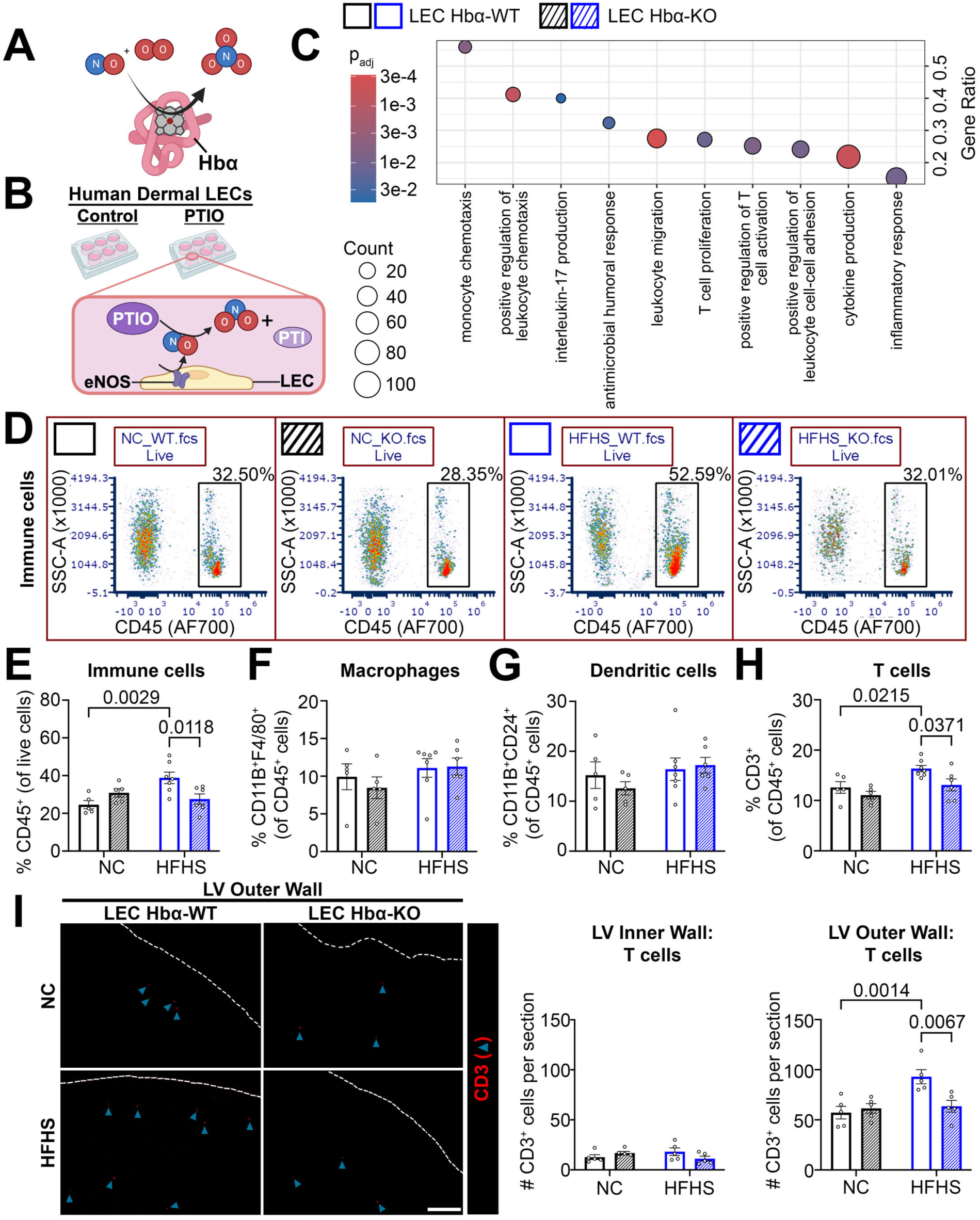
LEC Hbα promotes accumulation of T cells in the heart during CMD. (A) Graphic showing the nitric oxide (NO)-scavenging function of Hbα. (B) Experimental layout for bulk RNA-sequencing. Human dermal lymphatic endothelial cells (LECs) were treated with PTIO (100 μM, 24 hours) to mimic Hbα function in scavenging NO produced from endothelial NO synthase (eNOS). N = 3 biological replicates/condition. (C) Immune-related gene ontology terms enhanced in PTIO-treated LECs, identified from differentially expressed genes ranked by log2(FoldChange). (D) Representative density plots of cardiac immune cells from (NC)-fed and high fat, high sucrose (HFHS)-fed LEC Hbα-wildtype (WT) and -knockout (KO) mice. (E-H) Proportions of cardiac (E) immune cells, (F) macrophages, (G) dendritic cells, and (H) T cells, assessed using flow cytometry. NC LEC Hbα-WT and -KO groups: N=5 mice each; HFHS LEC Hbα-WT: N=7; HFHS LEC Hbα-KO: N=6. (I) Representative CD3 immunostains on the left ventricle (LV) outer wall, with wall-specific CD3^+^ T cell counts. White dashed line indicates start of LV outer wall. Blue arrows point to T cells. Counts totaled from 10-12 outer wall images and 4-5 inner wall images per heart. N=5 mice/group. Scale = 200 μm. Statistical significance determined by 2-way ANOVA with Šídák’s multiple comparisons test.

Through a vast lymphatic network, the heart is drained into mediastinal lymph nodes, which sample from the heart’s local environment to launch an immune response as needed. During T cell-mediated inflammation, T cell populations in the lymph node undergo clonal expansion following activation.^43^ To further confirm the regulation of local inflammation by LEC Hbα, we assessed T cell populations in mediastinal lymph nodes using flow cytometry (**Supplementary Figure 11A**). The proportions of total T cells, cytotoxic T cells, and T helper cells were all reduced in the mediastinal lymph nodes of HFHS-fed LEC Hbα-KO mice, suggesting a local suppression in T cell-mediated inflammation (**Supplementary Figures 11B-11E**). Conversely, LEC Hbα likely does not regulate systemic inflammation as HFHS-fed mice exhibited enlarged spleens relative to NC-fed controls independent of Hbα expression (**Supplementary Figures 11F-11G**). These data together implicate LEC Hbα as a driver of local T cell accumulation and inflammation in the heart during CMD.

### LEC Hbα promotes T cell adhesion to cardiac lymphatic vessels during CMD

To better define the mechanism through which LEC Hbα impacts T cells in the heart during CMD, we re-examined our gene set enrichment analysis (**Figure 4C**) and identified pathways related to T cell activation and immune cell adhesion as potential mechanisms by which LECs may regulate T cells (**Figures 5A-5B, Supplementary Figure 12A**). However, NO alone has also been shown to influence T cell behavior.^44^ Therefore, we first evaluated whether NO can directly regulate T cell activation. To do so, we activated Jurkat T lymphoblasts for 4 hours in the presence or absence of an NO donor, DETA-NONOate (100 μM), before assessment of CD69^+^ activated T cells using flow cytometry (**Figure 5C; Supplementary Figure 12B**). We found NO did not exhibit any regulatory role on T cell activation (**Figure 5C, Supplementary Figure 12C**). Co-culture of Jurkat T cells with LECs pre-treated with PTIO or DETA-NONOate similarly did not promote T cell activation (**Figure 5D, Supplementary Figure 12D**). These data were unsurprising as we did not observe positive transcriptional changes in genes encoding major histocompatibility complex (MHC) molecules, which are required for direct T cell activation, in PTIO-treated LECs (**Figure 5E**). In fact, some MHC I-encoding genes trended downwards or were significantly decreased with PTIO treatment and MHC II-encoding genes were undetected across the entire dataset. In line with these *in vitro* findings, we observed that LEC Hbα did not regulate the proportion of activated T cells, cytotoxic T cells, or T helper cells in hearts of mice with CMD, though there was elevated T cell activation generally in HFHS-fed mice relative to NC-fed controls (**Figure 5F, Supplementary Figures 10D, 12E-12F**).

**Figure 5.**
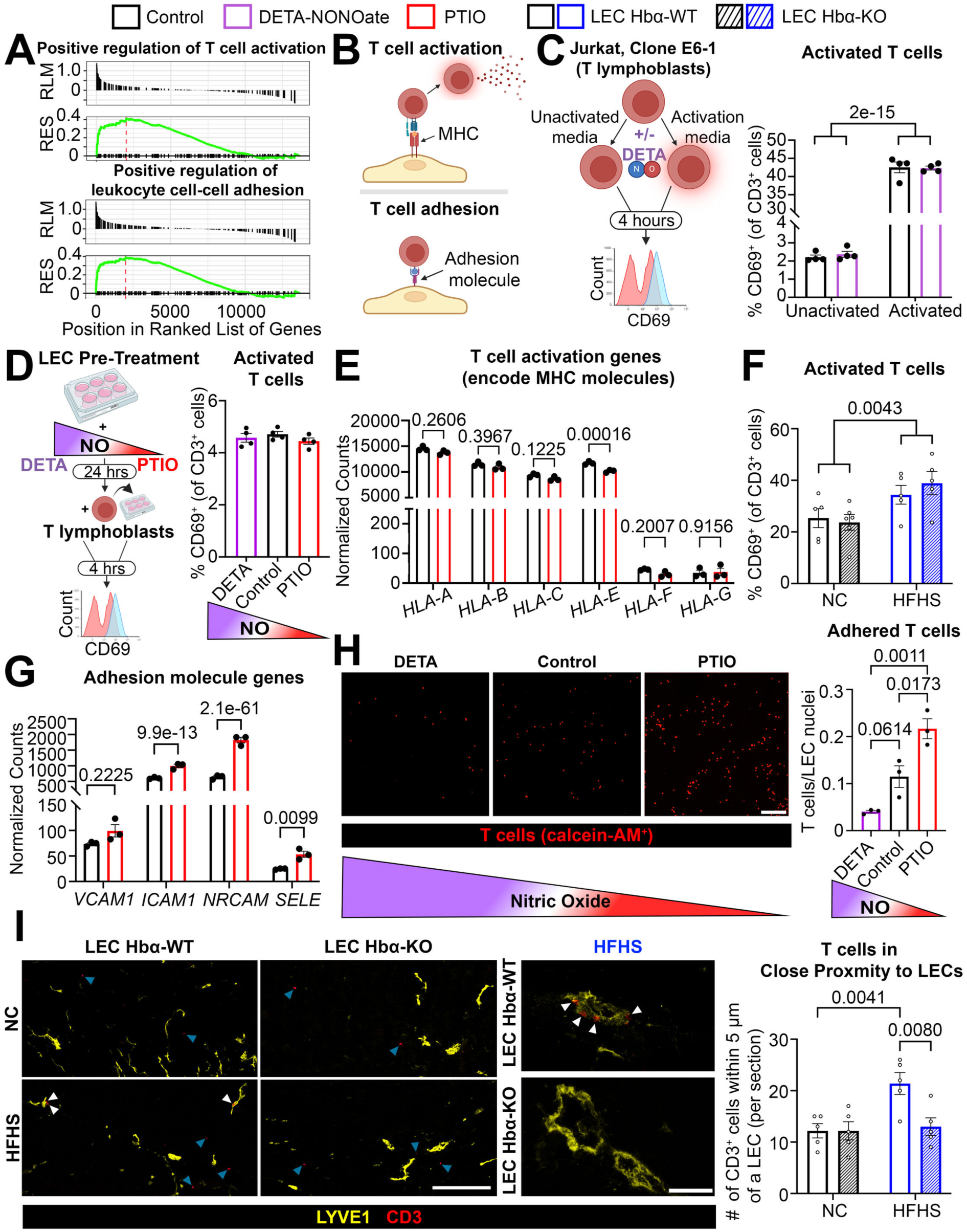
Local NO scavenging around LECs promotes T cell adhesion to LECs. (A) Ranked positions of genes from the gene sets “positive regulation of T cell activation” (top) and “positive regulation of leukocyte cell-cell adhesion” (bottom) in lymphatic endothelial cells (LECs) treated with the nitric oxide (NO) scavenger, PTIO (100 μM, 24 hours); genes ranked by log2(FoldChange) relative to control LECs. RLM = ranked list metric; RES = running enrichment score. (B) Scheme showing two different mechanisms by which LECs can regulate T cells: through activation (via antigen presentation by major histocompatibility complex [MHC]) and through adhesion. (C) *In vitro* experimental scheme. Jurkat T lymphoblasts were activated (50 ng/mL PMA + 0.1% ionomyocin, 4 hours) +/- the NO donor, DETA-NONOate (DETA; 100 μM). Activated (CD69^+^) T cells assessed using flow cytometry. N=4 biological replicates/group. (D) *In vitro* experimental scheme. Jurkat T lymphoblasts were co-cultured for 4 hours with LECs that were pre-treated with DETA- (100 μM), control-, or PTIO- (100 μM) media for 24 hours. Activated T cells assessed using flow cytometry. N=4 biological replicates/group. (E) Normalized counts from bulk RNA-sequencing for genes encoding MHC molecules in control and PTIO-treated LECs. N=3 biological replicates/group. (F) Flow cytometric analysis of activated T cells in hearts of normal chow (NC)- and high fat, high sucrose (HFHS)-fed LEC Hbα-wildtype (WT) and -knockout (KO) mice. NC LEC Hbα-KO: N=6 mice. All other groups: N=5 each. (G) Normalized counts from bulk RNA-sequencing for genes encoding adhesion molecules in control and PTIO-treated LECs. N=3 biological replicates/group. (H) Representative images of T cells (red) adhered to LECs (not pictured) pre-treated with DETA- (100 μM), control-, or PTIO- (100 μM) media for 24 hours, with quantifications on the right. N=3 biological replicates/group; 1 biological replicate = average of 3 technical replicates. Scale = 200 μm. (I) Representative immunostains for CD3 and LYVE1 in hearts from all groups, with quantification of T cells in close proximity to a LEC. N=5 mice/group. White arrows indicate T cells in close proximity to an LEC (< 5 μm from a LEC). Blue arrows indicate T cells > 5 μm from a LEC. Scale for lower magnification images = 200 μm; Higher magnification scale = 50 μm. Adjusted p-values in (E, G) determined using DESeq2. Statistical significance for other comparisons determined by (C, F, I) 2-way ANOVA with Šídák’s multiple comparisons test or (D, H) ordinary one-way ANOVA with Tukey’s post hoc analysis.

To determine whether LEC Hbα may instead promote T cell adhesion to LECs, we first checked for the expression of adhesion molecule genes in PTIO-treated LECs. While *VCAM1* expression was trending upwards with PTIO treatment, other documented adhesion molecule-encoding genes (*ICAM1*, *NRCAM*, *SELE*) were significantly increased relative to control LECs (**Figure 5G**). To validate these transcriptional changes, we performed an adhesion assay in which Jurkat T cells were co-cultured for one hour with LECs pre-treated with DETA-NONOate or PTIO (100 μM, 24 hours). This assay revealed that reducing NO bioavailability around LECs promoted T cell adhesion to LECs (**Figure 5H**). Immunostaining for both T cells and LECs in hearts from NC- and HFHS-fed LEC Hbα-WT and -KO mice corroborated this finding – HFHS-fed LEC Hbα-WT mice had significantly more T cells found in close proximity (< 5 μm) to a cardiac lymphatic vessel compared to NC-fed controls, and this was rescued in LEC Hbα-KO mice (**Figure 5I**). Taken together, these data suggest LEC Hbα promotes T cell adhesion to cardiac lymphatic vessels during CMD, which may explain the T cell accumulation observed during disease.

### Noncanonical NF-κB signaling is enhanced in LECs under conditions of reduced NO bioavailability

It is well described that nuclear factor kappa-light-chain-enhancer of activated B cells (NF-κB) signaling induces endothelial expression of immune cell adhesion molecules.^45^ Thus, we checked human bulk and single-nucleus RNA-sequencing from HFpEF and control hearts for NF-κB-encoding genes.^20, 33^ While transcript levels of the canonical NF-κB gene, *NFKB1*, were unchanged during HFpEF, the noncanonical gene, *NFKB2*, was significantly increased in HFpEF hearts (**Figure 6A**). Further, *NFKB2* transcripts trended upwards in HFpEF cardiac LECs (**Figure 6B**). In line with these data, human dermal LECs treated with PTIO had increased transcripts for *NFKB2* and *RELB*, which encodes the heterodimer protein involved in noncanonical NF-κB gene transcription (**Figure 6C**). *NFKB1* transcripts were modestly increased, though this difference was not statistically significant. To determine whether changes in NO bioavailability alter noncanonical NF-κB signaling in LECs, we immunoblotted for the *NFKB2* product, p100, and its mature form, p52, which is required for nuclear translocation and subsequent transcription of adhesion molecule genes (**Figures 6D-6E**). In agreement with the RNA-sequencing data on PTIO-treated LECs (**Figure 6C**), p100 protein levels were increased in PTIO-treated LECs relative to control (**Figure 6E**). There was also a trending increase in p52 levels in PTIO-treated LECs relative to DETA-NONOate-treated LECs (**Figure 6E**). Taken together, these data indicate noncanonical NF-κB signaling may be increased in LECs with NO scavenging, which could help to explain the pro-adhesive state of PTIO-treated and Hbα-expressing LECs.

**Figure 6.**
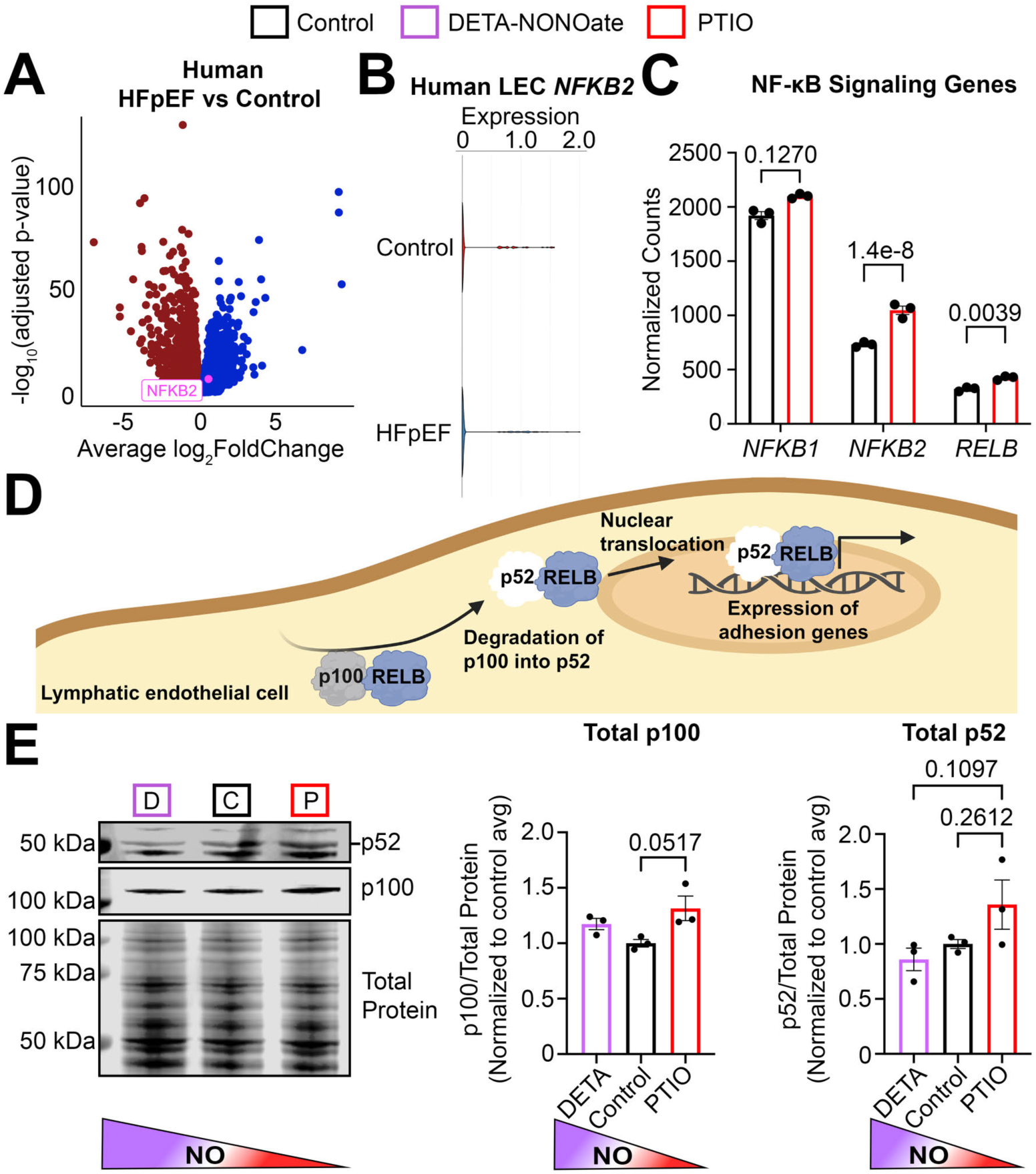
Noncanonical NF-κB signaling may be altered in LECs under conditions of altered NO bioavailability. (A) Differentially expressed genes from bulk RNA-sequencing of human heart failure with preserved ejection fraction (HFpEF) versus control hearts. Data re-analyzed from *Hahn et al*.^32^ (B) *NFKB2* expression in cardiac lymphatic endothelial cells (LECs) from control donors and HFpEF patients. Data re-analyzed from *Hahn et al*.^20^ (C) Normalized counts of genes encoding NF-κB signaling proteins from bulk RNA-sequencing of both control-treated LECs and LECs treated with the nitric oxide (NO) scavenger PTIO (100 μM) for 24 hours. N=3 biological replicates/group. (D) Graphic illustrating the processing of *NFKB2* product, p100, into p52 to promote nuclear translocation of p52 and subsequent transcription of adhesion molecule genes. (E) Total p100 and p52 from control-treated LECs as well as LECs treated with PTIO or the NO donor, DETA-NONOate (100 μM) for 24 hours; assessed by western blot. N=3 biological replicates/group. Adjusted p-values in (C) determined using DESeq2. Statistical significance for other comparisons determined by (E) ordinary one-way ANOVA with Tukey’s post hoc analysis.

### Loss of LEC eNOS in healthy mice mimics Hbα up-regulation in CMD

To further verify the crucial role of LEC-derived NO in the regulation of CMD, we generated *Flt4 Cre-ER^T2+/-^ Nos3^fl/fl^*(LEC eNOS-WT and -KO) mice to selectively delete eNOS from LECs. Male mice were administered tamoxifen to induce the genetic deletion and were then fed NC diet for 18 weeks, so that phenotypes observed could be directly linked to changes in LEC-derived NO (**Figure 7A**). Protein deletion was validated by loss of eNOS immunofluorescent signal in lymphatic vessels (**Figure 7B, Supplementary Figure 13A**). Importantly, arterial eNOS expression remained intact following tamoxifen induction (**Supplementary Figure 13B**). Body mass and spleen mass were both unchanged between LEC eNOS-WT and -KO mice (**Supplementary Figure 13C-13D**). In line with what was observed in mice with CMD (where Hbα was upregulated), eNOS deletion from LECs resulted in a reduction in cardiac lymphatic density only in the LV outer wall (**Figure 7C**). Cardiomyocyte hypertrophy was also observed in the LV outer walls of LEC eNOS-KO mice (**Figure 7D, Supplementary Figure 13E**). Importantly, the magnitude of cardiomyocyte area increase observed in the LV outer wall of LEC eNOS-KO mice was consistent with that of HFHS-fed mice with intact LEC Hbα expression (**Figure 7E**). The proportion of immune cells in the hearts of LEC eNOS-KO mice was unaltered (**Figure 7F**). However, the proportion of T cells in the heart was increased in LEC eNOS-KO mice, consistent with what we observed in Hbα-expressing mice with CMD (**Figure 7G**). Taken together, these data highlight the importance of LEC-derived NO in maintaining immune homeostasis and cardiac structural organization, while providing further support for the findings observed with Hbα upregulation in CMD.

**Figure 7.**
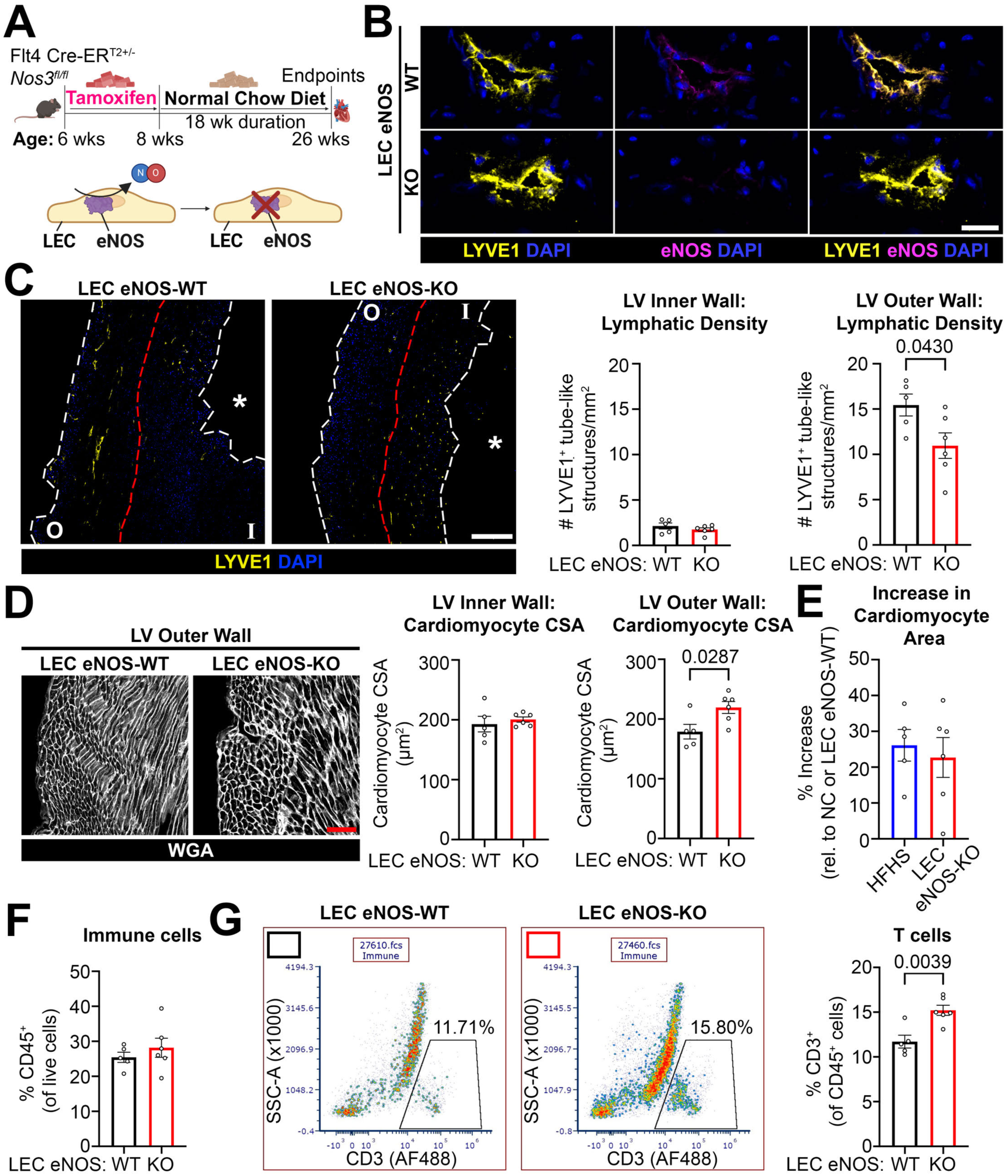
Loss of LEC eNOS in healthy mice promotes features of cardiometabolic disease. (A) Experimental scheme for Flt4 Cre-ER^T2+/-^ *Nos3^fl/fl^* mice. Endothelial nitric oxide synthase (eNOS) was deleted from lymphatic endothelial cells (LECs) to mimic Hbα up-regulation. LEC eNOS-wildtype (WT) and -knockout (KO) mice were then fed normal chow (NC) for 18 weeks (wks). (B) Representative LYVE1/eNOS co-stains on left ventricles (LV) from LEC eNOS-WT and -KO mice. Scale = 20 μm (C) Representative LYVE1 immunostains on LVs, with wall-specific quantifications of lymphatic density. LEC eNOS-WT: N=5 mice; LEC-eNOS-KO: N=6. * denotes LV chamber. O = outer wall, I = inner wall. Scale = 200 μm. (D) Representative wheat germ agglutinin (WGA) stains on LV outer walls, with wall-specific quantifications of cardiomyocyte cross-sectional area (CSA). LEC eNOS-WT: N=5 mice; LEC-eNOS-KO: N=6. Scale = 50 μm. (E) Percent increase in LV outer wall cardiomyocyte CSA in high fat, high sucrose (HFHS)-fed LEC Hbα-WT mice (relative to NC-fed controls) and LEC eNOS-KO mice (relative to LEC eNOS-WT controls). HFHS mice: N=5 mice; LEC eNOS-KO: N=6 mice. (F-G) Flow cytometric analysis of cardiac (F) immune cells and (G) T cells. LEC eNOS-WT: N=5 mice; LEC-eNOS-KO: N=6. Statistical significance determined by (C-G) unpaired t-test.

## Discussion

It is widely accepted that NO bioavailability is reduced during CMDs, such as HFpEF, though mechanistic work thus far has focused on reactive oxygen species and BECs as drivers of this phenomenon. In this work, we identified cardiac LECs as a key contributor to local reductions in NO signaling during CMD through expression of Hbα, thereby reproducing multiple features of the HFpEF phenotype, such as reduced cardiac lymphatic density, cardiac T cell accumulation, and hypertrophy development. Furthermore, we identified LECs as a crucial origin of NO production required to prevent the formation of CMD phenotypes in otherwise healthy mice.

Recent evidence highlighted hemoglobin beta as a potential modulator of nitrosative stress in both animal models of HFpEF and human patients.^12^ In this study, we identified genes encoding Hbα were also upregulated in murine and human HFpEF. In the human dataset^32^, *HBA1* and *HBA2* were the most highly up-regulated genes without any differential expression of genes specific to red blood cells. Thus, red blood cell contamination of HFpEF hearts can be ruled out for the interpretation of this finding. *In vivo* modeling of CMD revealed a 6-fold increase in the percentage of lymphatic vessels expressing Hbα. Yet, single-nuclei RNA-sequencing^20^ showed only a modest increase in *HBA2* expression and no expression of *HBA1* in LECs. With LECs making up only a small percentage of total cardiac cells, very few LECs were captured for analysis (N=148 control LECs; N=285 HFpEF LECs). Therefore, it is possible that a stronger increase in LEC *HBA2* expression could have been missed. BECs in murine CMD were also enriched with Hbα, but to a lesser magnitude than in LECs. Iron availability, myoendothelial junction formation, and transcription factors related to laminar shear stress have all been shown to regulate Hbα expression in BECs.^27, 29, 46^ However, it remains to be studied what is driving Hbα upregulation in pan-endothelial cells (and to a greater extent in LECs) during CMD.

LV hypertrophy is a common observation in patients with CMD.^4, 47^ We determined LEC Hbα promotes localized hypertrophic remodeling of cardiomyocytes to drive LV wall thickening and cardiac hypertrophy. Blood endothelial dysfunction is considered a driving factor of cardiac remodeling and dysfunction during CMD^2^, yet we observed no regulation of coronary vascular function by LEC Hbα. Furthermore, we did not observe changes in microvascular density under any condition. This was contrary to more severe models of CMD in which coronary microvascular rarefaction is commonly described.^31, 48, 49^ Relative to other models of CMD (e.g., the two-hit HFpEF model of high fat diet + L-NAME^31^), HFHS feeding produces a milder disease phenotype. HFHS-fed mice, along with maintaining coronary microvascular density, lack cardiac fibrosis and have a much milder hypertrophic response relative to mice undergoing the two-hit model. Thus, the findings of this study may be best applied in the context of early-stage CMD. Regardless, we were still able to recapitulate the reduction in cardiac lymphatic density in CMD that has been shown in both murine and human HFpEF alike.^19^ This suggests that perhaps lymphatic dysregulation precedes that of the blood vasculature during CMD progression, which is an entirely novel concept in the context of HFpEF. We further showed changes in cardiac lymphatic density to be dependent on LEC Hbα expression. As multiple studies have shown NO to be pro-lymphangiogenic^50, 51^, LEC Hbα likely represses lymphatic expansion directly through NO scavenging to account for the reduced cardiac lymphatic density observed in our CMD model.

Many others have shown NO bioavailability is reduced both systemically and locally in the heart during animal and/or human HFpEF.^7–12^ We recapitulated this in our CMD model, though LEC Hbα only contributed to local decreases in NO signaling. This was contrary to pan-endothelial Hbα expression, which we found regulated systemic NO levels. Other work has shown loss of eNOS from pan-endothelial cells reduced circulating nitrite levels, suggesting that BECs are key regulators of systemic NO bioavailability.^52^ From a therapeutic standpoint, this distinction is important to define as our findings indicate targeting of LEC Hbα alone would likely allow the persistence of systemic vascular dysfunction. Nitric oxide-based therapies, which have overall yielded mixed results, showed the greatest therapeutic promise based on their targeted effects on the systemic vasculature.^53^ Meanwhile, heart structure and function mostly remained unaltered. Based on therapy-specific limitations, the possibility remains that the cardiac lymphatic system was not properly being targeted in these trials to produce local benefit in the heart. Though, no trials have yet been published to probe this.

Through *in vitro* modeling, we determined local NO scavenging promotes an immunomodulatory transcriptional response in LECs. *In vivo* validation revealed LEC Hbα regulates immune cell accumulation in the heart during CMD. These findings align with classical anti-inflammatory properties associated with NO signaling.^54^ Other work has shown stimulation of cardiac lymphatic development in murine HFpEF completely prevents immune cell accumulation in the heart.^19^ Our work supports this finding and implicates LEC Hbα in the process. However, whether regulation of pan-immune cell accumulation during CMD occurs strictly from changes in lymphatic density mediated by Hbα versus being also from transcriptional changes in the LECs themselves remains to be shown.

During inflammation, the immune cells predominantly trafficked by lymphatic vessels include dendritic cells, macrophages, and T cells.^42^ Yet in CMD, we only observed LEC Hbα regulation of T cells. Perhaps Hbα expression modulates the expression of chemotactic and pro-adhesive signals which favor T cell regulation over other immune populations. It has also been shown in both humans and sheep that T cells make up ∼80-90% of immune cells in afferent lymph, which may make them more susceptible to accumulation in tissues when lymphatic dysfunction occurs.^55, 56^ The enrichment in T cells we observed in CMD hearts recapitulated findings reported in murine and human HFpEF.^20–22^ We found LEC Hbα drove this accumulation, which only occurred in the LV outer wall where lymphatics are localized. In murine HFpEF, T cells have been shown to promote cardiomyocyte hypertrophy.^21^ Nitric oxide signaling has also been shown to directly prevent cardiomyocyte hypertrophy.^15^ Thus, based on our findings alone, we are unable to definitively conclude whether LEC Hbα regulates cardiomyocyte hypertrophy directly via regulating local NO signaling or indirectly by promoting T cell accumulation in the heart.

T cell activation has recently been shown in human HFpEF hearts.^20^ While we also observed activated T cells in the hearts of mice with CMD, this was not regulated by LEC Hbα. To activate T cells directly, LECs must behave as antigen-presenting cells, yet we saw no difference in the expression of MHC-encoding genes with NO scavenging *in vitro*. Thus, T cell activation occurs through an Hbα-independent mechanism and/or different cell type during CMD. Aligning with the classical anti-adhesive roles of NO^54^, our *in vitro* studies revealed scavenging of NO surrounding LECs promoted a pro-adhesive state, with increased expression of adhesion molecule genes and increased T cell adhesion to LECs. Further, we determined there were increased T cells in close proximity to cardiac lymphatics vessels during CMD, but only with Hbα expression. Immune cell adhesion to LECs often precedes trafficking of the immune cell out of tissue.^42^ However, the overall enrichment of T cells in the heart during CMD when LECs express Hbα suggests instead either (1) adhesion to LECs is driving T cell retention in the heart or (2) lymphatic function is too impaired to properly clear adhered T cells. Lymphatic function in the heart is largely reliant on cardiomyocyte contraction.^57^ LV ejection fraction is unchanged in mice with CMD, indicating intact systolic function. Conversely, transcriptomic and *in vitro* data from LECs under conditions of reduced NO bioavailability showed up-regulation of *NFKB2* and a potential increase in non-canonical NF-κB signaling, which has been shown to induce adhesion molecules, including *VCAM1* and *ICAM1*, in other disease contexts.^58–60^ Thus, we postulate LEC Hbα is more so likely driving T cell accumulation in the heart during CMD through adhesion-based retention.

Our findings allowed us to identify LEC-derived NO as a crucial protective messenger during CMD development. We further confirmed this through generation of a novel LEC-specific eNOS-KO mouse, which exhibited reduced cardiac lymphatic density, cardiomyocyte hypertrophy, and cardiac T cell accumulation despite the absence of any disease stimulus. Contrary to what we observed in Hbα-expressing mice with CMD, LEC eNOS-KO mice did not have increased proportions of pan-immune cells in the heart. Perhaps genetic deletion of eNOS does not create a large enough inflammatory insult to observe a difference. This idea may be corroborated in part by the lack of splenomegaly in eNOS-KO mice. However, despite no expansion in total immune cells, the proportion of T cells in the heart was still increased. This further strengthens the argument that T cells are most susceptible to cardiac accumulation under conditions where LEC-derived NO is decreased, whether it be due to changes in lymphatic density alone and/or increased adhesion to LECs. These findings together define the necessity for LEC-derived NO in maintaining cardiac structural and immune homeostasis and highlight that loss of LEC-derived NO through eNOS deletion or Hbα upregulation promotes the development of a CMD-like phenotype.

## Novelty & Significance

### What is known?

- Nitric oxide bioavailability is reduced during cardiometabolic diseases, like heart failure with preserved ejection (HFpEF).
- Cardiac lymphatic vessels have reduced density and function during HFpEF.
- T cells accumulate in the HFpEF heart to modulate disease phenotypes, such as cardiac hypertrophy.

### What new information does this article contribute?

- Lymphatic endothelial cell-specific deletion of hemoglobin alpha (Hbα) prevents hypertrophic remodeling of the heart during cardiometabolic disease.
- Lymphatic endothelial cell-specific deletion of Hbα prevents T cell accumulation in the heart during cardiometabolic disease.
- Loss of endothelial nitric oxide synthase from lymphatic endothelial cells is sufficient to recapitulate features of cardiometabolic disease in otherwise healthy mice.

During cardiometabolic diseases, such as HFpEF, reduced nitric oxide bioavailability promotes hypertrophic remodeling of the left ventricle. Cardiac lymphatic vessel density is also reduced in this disease context. Here, we identified lymphatic endothelial cells, through increased expression of Hbα, are major contributors to reduced nitric oxide signaling locally in the heart during cardiometabolic disease. Loss of Hbα expression in lymphatic endothelial cells prevented cardiometabolic disease-induced hypertrophic remodeling in the heart. Lymphatic endothelial Hbα also regulated cardiac lymphatic density during cardiometabolic disease, without having any effects on the coronary blood vasculature. T cells have been implicated in driving cardiometabolic disease phenotypes. We found lymphatic vessels, through endothelial Hbα expression, promote T cell accumulation in the heart during cardiometabolic disease. Deletion of endothelial nitric oxide synthase from lymphatic endothelial cells in otherwise healthy mice recapitulated our findings from Hbα-expressing mice with cardiometabolic disease. Together, these findings novelly point to cardiac lymphatic endothelial cells as a crucial source of nitric oxide important for maintenance of both cardiac and immune homeostasis. Further, we identified Hbα as a novel regulator of nitric oxide signaling during cardiometabolic disease.

## Acknowledgments

The authors acknowledge the Research Histology Core, Flow Cytometry Core, Molecular Imaging Core, and Molecular Electron Microscopy Core at the University of Virginia School of Medicine for their expertise and resources. This work used the 9.4T MRI system in the Molecular Imaging Core Facility which is supported by the University of Virginia School of Medicine, Research Resource Identifiers (RRID): SCR_025472. Imaging data was acquired through the University of Virginia Molecular Imaging Core Lab, with S10OD025024: Bruker 9.4T BioSpec MRI small-animal scanner. Transmission electron micrographs were recorded using the 200kV FEI Tecnai F20 at the University of Virginia Molecular Electron Microscopy Core facility (RRID:SCR_019031), which is supported in part by the School of Medicine and built with NIH grant G20-RR31199.

## Non-standard Abbreviations and Acronyms

BEC: blood endothelial cell
CMD: cardiometabolic disease
eNOS: endothelial nitric oxide synthase
Hbα: hemoglobin alpha
HFHS: high fat, high sucrose
HFpEF: heart failure with preserved ejection fraction
HFrEF: heart failure with reduced ejection fraction
KO: knockout
LEC: lymphatic endothelial cell
LV: left ventricle
MHC: major histocompatibility complex
NC: normal chow
NF-κB: nuclear factor kappa-light-chain-enhancer of activated B cells
NO: nitric oxide
PLN: phospholamban
WT: wildtype

